# Cross-Lagged Panel Model in Medical Research: A Cautionary Note

**DOI:** 10.1101/486217

**Authors:** Satoshi Usami, Naoya Todo, Kou Murayama

## Abstract

Longitudinal designs provide a strong inferential basis for uncovering reciprocal effects or causality between variables. For this analytic purpose, a cross-lagged panel model (CLPM) has been widely used in medical research, but the use of the CLPM has recently been criticized in methodological literature because parameter estimates in the CLPM conflate between-person and within-person processes. The aim of this study is to present some alternative models of the CLPM that can be used to examine reciprocal effects, and to illustrate potential consequences of ignoring the issue. A literature search, case studies, and simulation studies are used for this. We examined more than 300 medical papers published since 2009 that applied cross-lagged longitudinal models, finding that in all studies only a single model (typically, the CLPM) was performed and potential alternative models were not considered to test reciprocal effects. In 49% of the studies, only two time points were used, which makes it impossible to test such alternative models. Case studies and simulation studies showed that the CLPM often has worse model fit and markedly different estimates of cross-lagged parameters than alternative models, suggesting that research that relies on the CLPM only may draw erroneous conclusions regarding the presence, predominance, and sign of reciprocal effects as well as about causality.

## Cross-Lagged Panel Model in Medical Research: A Cautionary Note

Collecting longitudinal data has become widely popular in medical research and other disciplines due to its statistical advantages over cross-sectional data. One of the biggest advantages of using a longitudinal design is that it can provide richer information for statistical inference aimed at uncovering reciprocal effects or causality between variables to answer questions such as how change (or growth, development) in one variable affects that of the other. More than 30 years ago, Nesselroade and Baltes^1^ reviewed the benefits and drawbacks of using longitudinal data in psychology, noting that revealing causes (determinants) of intra-individual change is one of the major strengths of longitudinal data. Likewise, in the econometrics literature, Hsiao^2^ argued that panel (i.e., longitudinal) data is effective for inferring dynamic relations between variables.

One of the most common methods for addressing reciprocal effects in medical research is use of a cross-lagged panel model (CLPM; Duncan^3^; also known as a dynamic panel model, autoregressive cross-lagged model, cross-lagged path model, or cross-lagged regression model), especially after the CLPM was integrated into the framework of structural equation modeling (e.g., Finkel^4^, Marsh and Yeung^5^). In these models, reciprocal effects are examined by testing the cross-lagged relations, which are the effect of variable *X* on variable *Y* after controlling for the previous effects of *X*.

The CLPM is a simple and powerful model to test reciprocal effects, and thus it has been widely used. However, the application of the CLPM has also recently been criticized. Notably, Hamaker, Kruiper, and Grasman^6^ criticized the use of the CLPM because the cross-lagged estimates in the CLPM conflate between-person and within-person processes, and so the results do not represent the actual within-person relations over time. Between-person relations are the covariation of two variables in terms of individual differences (e.g., individuals with higher *X* tend to have higher *Y* relative to individuals with lower *X*), whereas within-person relation are the covariation within one person of two variables across time points or situations. Obviously, these two types of relations are conceptually different. As such, the fact that estimates from traditional CLPM conflate between-person and within-person relations means that the cross-lagged estimates from the CLPM are conceptually difficult to interpret. Indeed, the importance of disaggregation to examine within-person processes has been widely acknowledged in the methodological literature (Curran & Bauer^7^; Hamaker^8^; Hoffman & Stawski^9^). Relying on the CLPM may draw erroneous conclusions regarding the presence, predominance, and sign of reciprocal effects as well as about causality.

To address this inherent problem with the CLPM, Hamaker et al^6^ proposed a random-intercepts CLPM (RI-CLPM) as a possible analytic option. As discussed later, in the RI-CLPM, individual differences are effectively controlled by the inclusion of a latent variable that represents a time-invariant (but person-variant) trait-like factor; this allows testing the reciprocal effects within individuals. If this model is extended to include measurement errors, the model is equivalent to a so-called (bivariate) stable trait autoregressive trait and state (STARTS) model (Kenny & Zautra^10, 11^). Usami, Murayama, and Hamaker^12^ discussed the mathematical and conceptual relations between various cross-lagged models, including these models.

These recent studies are insightful and informative, providing applied medical researchers a basis for thinking about how to test reciprocal relations by longitudinal data. However, the arguments are limited mostly to mathematical and conceptual relations. As a result, we still know little about whether, when, and how the choice of different cross-lagged longitudinal models has substantive consequences for parameter estimates of reciprocal effects in practice, leading researchers to draw different conclusions from the same data in medical sciences. The aim of the current manuscript is to show the practical implications and importance of considering these alternative models when investigating reciprocal effects. This is approached through a literature search, case studies, and statistical simulations. In the literature search, we first investigate the current common practice of longitudinal research in the medical literature, showing that medical researchers rely heavily and almost exclusively on the traditional CLPM when testing reciprocal relations or causality, and do not consider potential alternative models. Then, with case studies and statistical simulations, we illustrate the potential danger of this common practice, showing it can result in mistaken conclusions about reciprocal effects. In the end, we also provide some practical guidelines, hoping to help applied medical researchers who work on longitudinal data in the future.

## Cross-Lagged Longitudinal Models

In this paper, we focus on three cross-lagged longitudinal models: the (traditional) CLPM, the RI-CLPM, and the STARTS model. Below, following Usami et al, ^12^ we describe these models by emphasizing the commonalities and differences among these cross-lagged models. Throughout the paper, we assume that researchers are interested in the reciprocal relation between two variables *X* and *Y*, although it is easy to expand the models in a way that include more than two variables (e.g., when examining mediating effects of variables is a main focus of the research).

### CLPM

Let *x*_*it*_ and *y*_*it*_ be the measurements at time point *t* (1…*t*…*T*) for individual *i* (1…*i*…*N*). In the CLPM, *x*_*it*_ and *y*_*it*_ are first modeled as

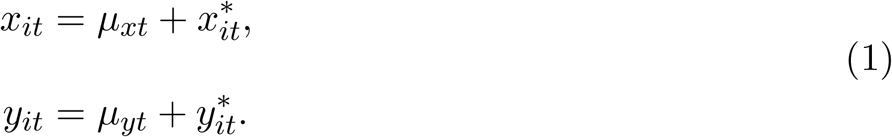

Here *μ*_*xt*_ and *μ*_*yt*_ are the temporal group means at time point *t*; 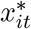 and 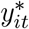 are temporal deviation terms from the temporal group means for individual *i*. With these equations, the trajectories of the temporal group mean are implicitly removed from the raw data. By definition, the deviations have a mean of zero. Then, *x*_*it*_ and *y*_*it*_ for *t* ≥ 2 are modeled as

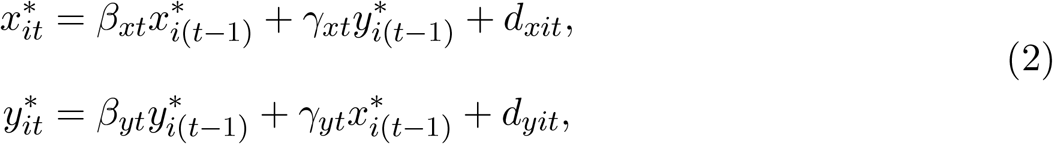

where *β*_*xt*_ and *β*_*yt*_ are autoregressive parameters and *γ*_*xt*_ and *γ*_*yt*_ are cross-lagged regression parameters at time point *t*. For these parameters, time-invariance can also be assumed (by using *β*_*x*_ and *β*_*y*_, and *γ*_*x*_ and *γ*_*y*_) if the cross-lagged relationships are assumed to be stable over time. Note that with *t* = 1, the initial observations *x*_*i*1_ and *y*_*i*1_ are modeled as exogenous variables.

From the view of Granger causality (Granger^13^), estimates of cross-lagged regression parameters (the longitudinal relationship between *Y*_*t*–1_ and *X*_*t*_ after controlling for the baseline *X*_*t*–1_) are key for inferring reciprocal relations between the variables. The residuals *d*_*xit*_ and *d*_*yit*_ are usually assumed to be normally distributed and correlated:

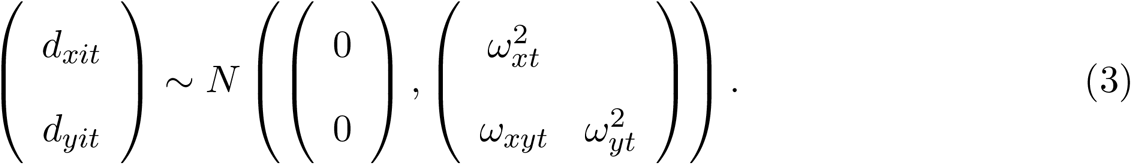

Here, 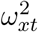 and 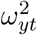 are time-variant residual variances and *ω*_*xyt*_ is a time-variant residual covariance. As with previous parameters, time-invariant residual variances and covariances can also be assumed (by using 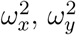, and *ω*_*xy*_). A path diagram of the CLPM is provided in Figure 1a.

### RI-CLPM

In the RI-CLPM (Hamaker et al^6^), *x*_*it*_ and *y*_*it*_ are modeled as

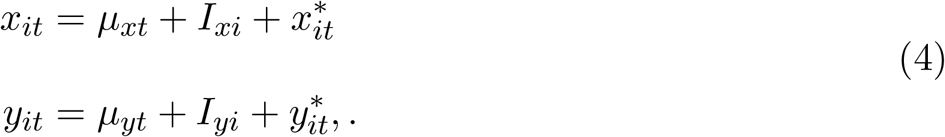

Again, *μ*_*xt*_ and *μ*_*yt*_ are the temporal group means. Critically, the model also includes *I*_*xi*_ and *I*_*yi*_, which are the defining characteristic of the RI-CLPM. These are (time-invariant) trait factors that represent individual’s trait-like deviations from temporal group means. Trait factors *I*_*xi*_ and *I*_*yi*_ have means of 0 and variance-covariance matrix ***V***. By accounting for trait factor scores, for each individual, 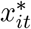 and 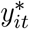 represent temporal deviations from the means of that individual because they are subtracted from the expected scores of individual *i* (i.e., *μ*_*xt*_ + *I*_*xi*_ and *μ*_*yt*_ + *I*_**yi**_). Accordingly, in the RI-CLPM, the time series 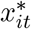 and 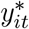 can be considered as within-person fluctuation. Due to this statistical property in temporal deviations, at *t* = 1 the initial deviation terms (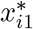 and 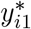) are assumed to be uncorrelated with the trait factors. Using these within-person deviation terms, in the RI-CLPM the cross-lagged relations are modeled as in the Equation 2 for *t* ≥ 2. A path diagram of the RI-CLPM is provided in Figure 1b.

Because the RI-CLPM accounts for trait factors and then separates stable between-person differences (i.e., trait factors) from within-person fluctuations over time, cross-lagged relations in the RI-CLPM can be considered as the one pertaining to a process that takes place at the within-person level. Therefore, in the RI-CLPM, *γ*_*x*_ and *γ*_*y*_ can be interpreted as the quantity that express the extent to which the two variables influence each other *within* individuals. Because longitudinal data typically include both quantitative information of within-person changes and its individual differences, the CLPM, which does not account for trait factors (i.e., individual differences), fails to disaggregate these two components. As such, the CLPM provides inaccurate estimates for within-person reciprocal effects.

Note that if substituting the cross-lagged relations of Equation 2 into 4, the trait factors, which are separated from independent variables (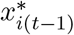 and 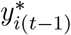), can obviously be interpreted as random intercepts in the model. The model is named after this statistical fact. Obviously, the CLPM is a special case of the RI-CLPM, found by letting *I*_*xi*_ = 0 and *I*_*yi*_ = 0. The RI-CLPM requires two or more variables to have been measured at three or more time points, while the CLPM requires only two time points.

### STARTS model

By extending the RI-CLPM to include measurement error, we obtain the STARTS model (Kenny & Zautra^10, 11^). In the (bivariate) STARTS model, *y*_*it*_ and *x*_*it*_ are decomposed into latent true scores *f*_*xit*_ and *f*_*yit*_ and measurement errors *ϵ*_*xit*_ and *ϵ*_*yit*_.

That is,

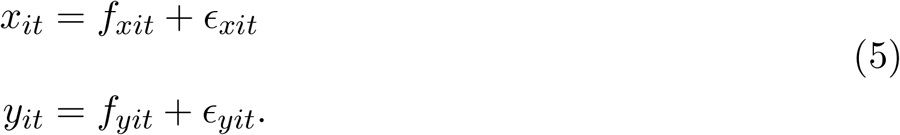

These measurement errors are usually assumed to be normally distributed and possibly correlated, that is,

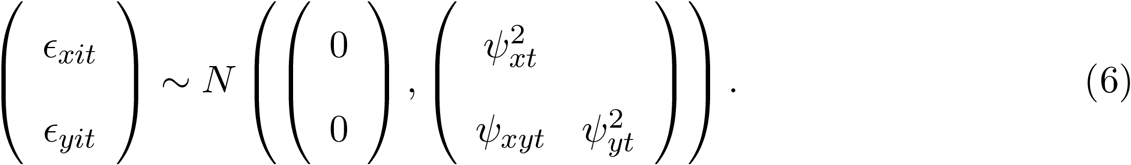

Here, 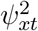 and 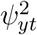 are measurement error variances, and *ψ*_*xyt*_ is an error covariance. If needed, time-invariant measurement error (co)variances can be assumed. As in the RI-CLPM, *f*_*xit*_ and *f*_*yit*_ are modeled as

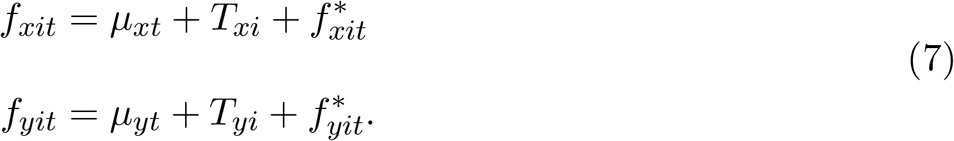

Here, 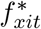 and 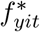 are the terms expressing temporal deviation from the expected scores of individual *i*, with accounting for measurement error.

Substituting the equation 7 into the equation 5 provides the specification of the STARTS model:

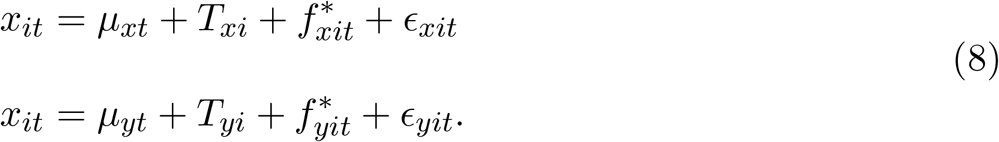

As in Eq. 2, temporal deviation terms are modeled as

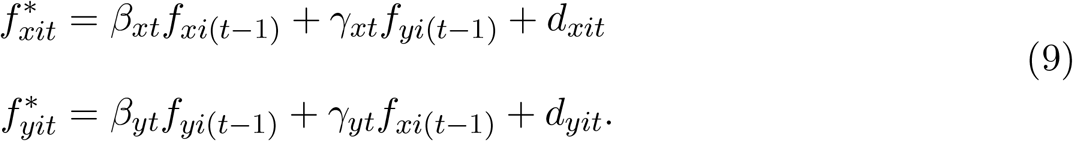

A path diagram of the STARTS model is provided in Figure 1c. Obviously, in the STARTS model, cross-lagged relations are posited between latent true scores, rather than between observed scores, distinguishing it from the RI-CLPM and the CLPM. However, the STARTS model and the RI-CLPM share a common critical feature—the inclusion of trait factors. As such, like the RI-CLPM, cross-lagged parameters (*γ*_*xt*_ and *γ*_*yt*_) in the STARTS model reflect within-person reciprocal effects. The STARTS model requires two or more variables to have been measured at four or more time points. This means that we can compare RI-CLPM and the STARTS to determine which of these models fits better to the data so long as more than three waves are available.

When observations may be influenced by measurement errors occurring for procedural reasons, accounting for measurement errors is desirable. However, the specification of measurement error when there is only one indicator variable (such as in the STARTS model) sometimes involves costs in terms of parameter estimation. Indeed, research has reported that the STARTS model often encounters estimation problems such as improper solutions and non-convergence. Conceptually, one primary reason is the fact that unlike trait factor variances (*ν*^2^) and residual variances 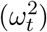, the contribution from measurement error variances 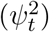 is temporal: in the model-implied variance-covariance matrix, 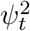 appears at time point *t* only. Because of this, unstable estimates of some parameters (particularly autoregressive parameters) caused by some aspects of the research design (e.g., small sample size) can easily inflate the variances of the deviation terms 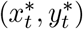, increasing the risk of obtaining negative estimates of 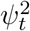.

Therefore, previous studies have also proposed models that incorporate multiple indicators (rather than a single indicator) to represent latent variables (see Cole et al^14^; Luhmann, Schimmack, & Eid^15^). In addition, research has also suggested the utility of a Bayesian approach to avoid unstable parameter estimation (Lüdtke, Robitzsch, & Wagner^16^).

## Review of the literature

### Method

To investigate recent trends in the use of cross-lagged longitudinal models in medical research, we conducted a literature search through the UTokyo REsource Explorer (TREE; http://tokyo.summon.serialssolutions.com/) web search engine in June of 2017. TREE aggregates information from many major databases (e.g., Web of Science, PubMed, PsycINFO, Engineering Village, ERIC, JSTOR) and electronic journals under contract with The University of Tokyo. TREE summarizes this collection of information in a single search window, allowing us to perform more comprehensive and efficient literature search than by using the individual databases separately. We first used the English keywords “cross lagged model” and “cross lagged relation”, searching English papers published since 2009 in medical journals. In addition, we limited our search to only peer-reviewed papers. Therefore, news items, book reviews, and doctoral dissertations were not considered.

We found 324 medical papers by this method. Of these, we excluded 53 papers that did not apply any cross-lagged longitudinal models to actual data, leaving us with 271 papers. Most of the excluded papers were review papers, statistical simulations, or methodological and statistical discussion. See Table 1 for the complete list of retained papers.

**Table 1.**
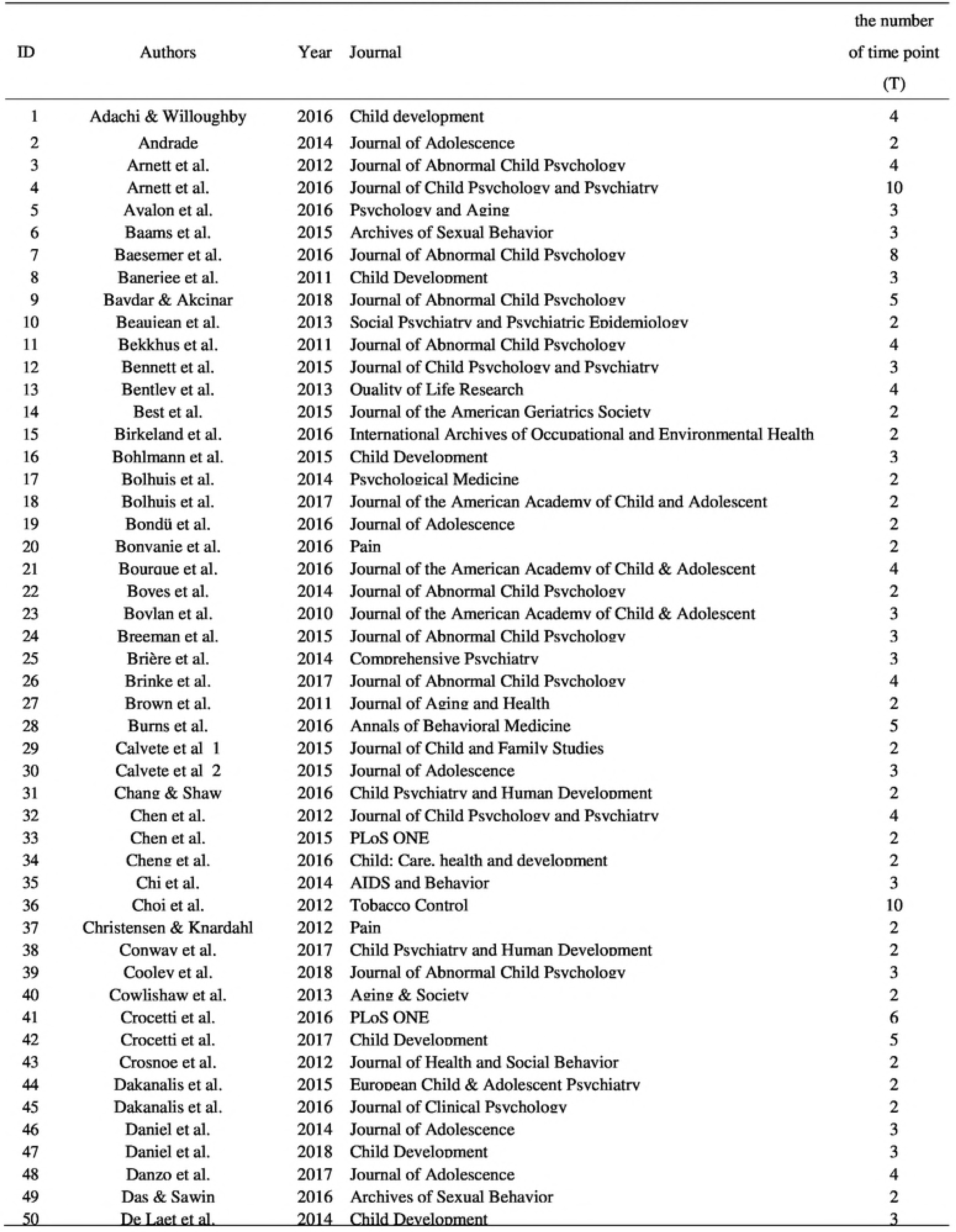
The list of 271 papers that applied cross-lagged models.

### Result

Among 271 papers, 106 (= 40%) papers collected longitudinal data at two time points. 89 (= 33%) papers collected data with three waves, 36 (= 13%) with four waves, 16 (= 6%) papers with five waves, and 24 (= 9%) at more than five time points. The = 40% proportion for two time points is close to the one = 45% reported by Hamaker et al^6^ in the field of psychology. With regard to the statistical analysis they performed, 257 papers (= 95%) used the CLPM to analyze longitudinal data, and one paper used a model similar to the RI-CLPM (see Telley et al, 2015 in Table 1; this model does not assume autoregressive parameters). Other papers applied different models, such as an autoregressive latent trajectory model (Poirier et al, 2016), a latent change score model (LCS; Baydar and Akcinar, 2018; Natsukai et al, 2013; Occhipinti et al, 2015; Usami et al, 2015), a model similar to the latent curve model with structured residuals (Baams et al, 2015; Mustillo et al, 2012; Williams et al, 2011), or a fixed-effects regression model (Baesemer et al, 2016; a model similar to the LCS). For the mathematical and conceptual relations between these models, see Usami et al.^12^ Five papers used a multilevel-model framework (Arnett et al, 2016; Cooley et al, 2018; Daniel et al, 2018; Fuller-Tyszkiewicz et al, 2015; Kashdan et al, 2014) to account for individual differences in parameters of the cross-lagged model (see General Discussion on this point). Note that no research applied the STARTS model, and few studies compared analysis results from different cross-lagged models (one exception is a methodological paper of Usami et al, 2015, which compared analysis results from the LCS model and the CLPM).

These results indicate the heavy reliance on the traditional CLPM in the literature. It is also important to note that alternative cross-lagged longitudinal models (e.g., the RI-CLPM and the STARTS model) require at least three time points (with a stability assumption; the STARTS model requires at least four time points with an instability assumption) to fit the model (for the ALT model, we need four time points with a stability assumption). The fact that about 40% of the papers collected data at only two time points suggests that almost half of applied medical research implicitly precludes the option of using these alternative models.

## Case studies

### Method

To compare analysis results based on different cross-lagged longitudinal models, we focused on the 165 papers that collected longitudinal data with more than two time points. Among them, we randomly selected 50 papers and using the contact information provided in each of the paper we contacted the corresponding authors of the papers via email to request they share the dataset to help our research. In this contact, we emphasized that (1) our primary research purpose is simply to compare analysis results from different cross-lagged models, not to criticize their findings, (2) we would not provide any estimation results from the original paper or relevant information in the datasets to prevent identification of the source of the paper, (3) we would not share the dataset with any other researchers, and that (4) we did not need information about variables that are not relevant to cross-lagged analysis (e.g., personal information of participants).

To increase response rates from authors, we contacted the authors after one month if we had not received a reply from the first contact. As a result, we received a total of 21 responses from the authors (response rate: 42%), and among them, five authors (from five different papers) granted us access to their datasets. We were unable to obtain permissions from the authors of the other 16 papers, mainly because sharing with us might have violated the data sharing policy of their sources. Among the five datasets, two datasets were publicly available online without special permission from the authors, two datasets were provided directly by the authors, and one dataset was provided after a review of the data use agreement that we submitted. Note that one of the datasets provides us with the access only to the sample means and sample (co)variances information (rather than the raw data), which allowed us to estimate the parameters but not to fully account for missing data.

Among five datasets, two datasets have three time points and the others have more than three time points (*M*_time–points_ = 6.0). The average sample size of these datasets is large (*N* = 2,741). In this paper, we do not give the exact number of participants and time points for each study to prevent the identification of the studies. While all five studies applied CLPM, some of them specified the model in slightly different ways. Specifically, two studies assumed second-order autoregressive and cross-lagged parameters as well as first-order parameters. Another study assumed a mediator between two variables. In addition, one study assumed time-invariant parameters (i.e., stability), while the other four studies did not.

To ensure the comparability of the results between datasets, in the current analysis, we assume time-invariant parameters for autoregressive and cross-lagged coefficients (*β* and *γ*) and residual and error (co)variances (*ω*^2^ and *ψ*^2^). In addition, neither second-order parameters nor external variables (e.g., mediators) were included in any of the analyses. This setup also means that the results reported in the current paper are all different from those reported in the original papers. Note that one study collected multi-group data and applied the CLPM using multi-group analysis. For this dataset, we assumed group-invariant parameters for autoregressive and cross-lagged coefficients as well as residual and error (co)variances (i.e., measurement invariance between groups) while setting no constraints on the difference of temporal means between groups.

All analyses were conducted using Mplus version 7.4 (Muthen & Muthen^17^). However, we found improper solutions and non-convergence in four of the five datasets when using maximum likelihood (ML) estimation to fit the RI-CLPM or the STARTS model. In such cases, we instead used Bayes estimation, based on a Markov chain Monte Carlo method under the assumption of non-informative priors. With Bayes estimation, we obtained parameter estimates successfully without any convergence problems. For more detailed discussion about ML and Bayes estimation in terms of estimation problems in applying the STARTS model, see Liidtke, Robitzsch, & Wagner^16^.

### Result

Table 2 provides (unstandardized) autoregressive/cross-lagged parameter estimates and standard errors for the CLPM, the RI-CLPM, and the STARTS model. Except for the cross-lagged parameter estimates in Research 2, all autoregressive/cross-lagged parameter estimates with the CLPM were statistically significant with two-sided *α* = .05. This can be partly attributed to the large sample sizes in these datasets, which increased the statistical power.

Although the RI-CLPM and the STARTS model also showed significant estimates in most cases, 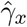 is not statistically significant in Research 4, while it is significant with the CLPM. Another different result is that the sign of 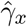 in the STARTS model was different from that with the CLPM in Research 3.

We also found notable differences in the magnitudes of parameter estimates among cross-lagged models. The RI-CLPM provided smaller autoregressive parameter estimates 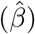 than the CLPM did (approximately 0.49 times the size), while the STARTS model provided larger estimates on average (approximately 1.45 times the size).

The relation between parameter estimates from different cross-lagged longitudinal models must depend in complicated ways on the magnitude of the parameter values and on research design factors (e.g., *N* and *T*), and we need to be careful when generalizing the findings. But, one potential explanation for the increased autoregressive parameters in the STARTS model is the dissociation of measurement errors in the model because the autoregressive parameters are the major source of correlations (i.e., the variance-covariance matrix) between time points. For the RI-CLPM, in contrast, the decreased autoregressive parameter estimates may be a consequence of trait factors, which would explain a large portion of the correlations between time points.

The differences in estimates of autoregressive parameters between the RI-CLPM and the STARTS model also lead to differences between their cross-lagged parameter estimates and those found by the CLPM. In this case study, the RI-CLPM and the STARTS model showed smaller cross-lagged estimates (in absolute value, 0.66 and 0.62 times the size, respectively) from those with the CLPM. Although we need to be careful about the generalizability of findings, it is well-known that the magnitude of within-cluster (in this case, within-person) relations (i.e., cross-lagged parameters in the RI-CLPM and the STARTS model) is smaller than those of between-cluster (in this case, between-person) relations, when the between-cluster difference is larger than the within-cluster difference. The decreased cross-lagged effects could be explained by this so-called ecological fallacy (Robinson^18^).

With regard to standard errors, interestingly, the standard errors of 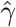 in the RI-CLPM and the STARTS model are, on average, 1.6 and 2.7 times, respectively, the size of those with the CLPM. These results indicate that the inclusion of parameters that are specific to these models (i.e., trait factor (co)variances in the RI-CLPM and those and error (co)variances in the STARTS model) leads to an increase in standard errors. In combination with the observed upward or downward changes in autoregressive and cross-lagged parameter estimates, these results indicate that the RI-CLPM and the STARTS model will produce substantially different results on statistical tests than the CLPM will.

It is also important to note that, among the five datasets, the CLPM was chosen as the best model in terms of model fit only once, when the Bayesian Information Criterion was used in Research 2. This result indicates that many previous studies that applied only the CLPM may have drawn erroneous conclusions about the magnitude and presence of reciprocal effects.

The results described here indicate the importance of comparing alternative models when testing for reciprocal effects, and the potential (in most cases, unintended) consequences of not considering multiple models. However, one might be concerned about the generalizability of the results due to the small number of studies (i.e., five) presented here. Another important issue is the improper solutions observed in two of the five datasets when applying the STARTS model. To address these issues more extensively, we conducted two statistical simulation studies, one focusing on the frequency of improper solutions and the other focusing on parameter estimates. Although the previous case studies indicated that these models could produce overtly different parameter estimates, to the best of our knowledge, no previous research has performed statistical simulation that directly compared the parameter estimates (and associated standard errors) produced by different cross-lagged longitudinal models we discussed here (i.e., the CLPM, the RI-CLPM, and the STARTS model). In addition, although some past studies have examined the frequency of improper solutions, focusing especially on the STARTS model (e.g., Cole et al^14^; Lüdtke et al^16^), no studies have systematically investigated the differences of longitudinal models used and examined the potential impact of model misspecification. Our statistical simulation also aims to extend the previous studies by addressing these points.

## Simulation Study

### Frequency of improper solutions

To systematically investigate the rate of improper solutions under various conditions, we performed Monte Carlo simulations, where both data generation model and analysis models were selected from the three models we have discussed, resulting in 9 (= 3 × 3) combinations of data generation and analysis models. This way, we can examine the potential influence of model misspecification (as well as the correct model specification) on improper solutions. For simplicity of the simulations, the stability of parameters was assumed.

For data generation, we systematically changed the number of total participants (*N* = 200, 600, 1, 000), the number of time points (*T* = 4, 6, 8), and the size of autoregressive parameters (*β* = *β*_*x*_ = *β*_*y*_ = 0.5, 0.7, 0.9). In this simulation, cross-lagged parameters γ were all fixed to 0.2. For the STARTS model, measurement error variances were set to 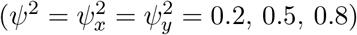. For the other models, *ψ*^2^ is always set to zero. Variances of the temporal deviation terms at the first time point (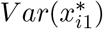 and 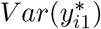), which are equivalent to those of observations in case of the CLPM, were fixed to 1 - *ψ*^2^. The size of *β* reflects the determination coefficients in cross-lagged regressions. For models with trait factors (i.e., the RI-CLPM and the STARTS model), we posited normal distribution for the trait factors and their variances were set to the half size of those of temporal deviation terms at the first time point (i.e., to 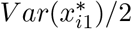 and 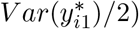.

Without loss of generality, the temporal group means were set to *μ*_*xt*_ = *μ*_*yt*_ = *t* – 1 for each time point. Correlation of the trait factors was set to 0.2. Correlation of temporal deviation terms at the first time point was set to 0.2, and in the STARTS model (time-invariant) correlations between measurement errors were set to 0.2. Finally, residual variances were fixed to 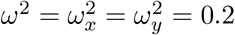, and correlation of residuals between variables was fixed to 0.2 for each time point.

We generated simulated data (200 trials for each combination) by crossing these factors, resulting in 81 (= 3(*N*) × 3(*T*) × 3(*β*) × 3(*ψ*^2^)) combinations of factors for each pair of data generation model and data analysis model. Each simulated dataset was analyzed by the three types of analysis models, and we counted the number of improper solutions, which was defined as (1) out-of-range parameter estimates (e.g., negative variances parameters) or (2) a singular approximate Hessian matrix after termination of iteration. The whole simulation procedure, including data generation and analysis, was conducted in R (R Core Team^19^) using the lavaan (Rosseel^20^) package with the ML estimation method. Simulation code is available in the Online Supporting Materials.

Table 3 presents the marginal proportions of improper solutions observed with each data analysis model under each level of the factors we manipulated. When the CLPM is used for analysis, it did not show improper solutions under any conditions. When CLPM is used for data generation, Table 3 shows that RI-CLPM and the STARTS model showed very large proportions of improper solutions (in the range of 40%–100%). Notably, in cases of the STARTS model, which posited measurement error (co)variances and residuals, 90% of the results exhibited improper solutions. Interestingly, the manipulated factors, such as the number of total participants (*N*) and number of time points (*T*) did not influence the results much. These results indicate that the impact of model misspecification dominates the risk of improper solutions, with the factors being manipulated playing a much smaller role. The same pattern was observed with different data generation models. Model misspecification was the biggest cause of improper solutions, and the STARTS model especially produced a higher number of improper solutions.

One particularly important observation is that improper solutions were still observed in the STARTS model even when the model was correctly specified. Indeed, the proportion of improper solutions was unacceptably high, at more than 70%. Note that, even compared with previous investigations (Cole et al^14^; Lüdtke et al^16^), our simulations showed larger number of improper solutions. This might be attributed to differences in the stability of measurements between the current simulations and the simulations in the previous studies. Instead of controlling the residual variances, the variances of all variables were set to 1 in the simulations of both Cole et al^14^ and Ludtke et al, ^16^, while we did not do this in the current investigation. In most of the current simulation conditions, the variances of variables are implicitly assumed to increase over time, as is often the case with longitudinal data in developmental/clinical research. Thus, the relative impacts of trait factor variances, (time-invariant) measurement error variances, and residual variances on observations become smaller at later time points, increasing the risk of out-of-range estimates in these variance estimates. Another important difference is that such previous investigations have considered univariate (rather than bivariate) version of the STARTS model. The bivariate version of the STARTS model, which we simulated in the current study, might have a bigger risk of improper solutions caused by a singular Hessian matrix.

For correctly specified models, the RI-CLPM performed better, especially when sample size and the number of time points were larger. However, the proportion of improper solutions was still not negligible (at 10– –15%). Therefore, although the RI-CLPM and the STARTS model can be considered as alternatives to the CLPM when investigating within-person reciprocal relations, these models might be susceptible to improper solutions, especially in the presence of model misspecification.

### Statistical properties of estimates

To investigate the statistical properties of cross-lagged parameter estimates in each cross-lagged longitudinal model, we performed another Monte Carlo simulation. As in the previous simulation, the data generation model and analysis model were selected from the three types of models. For data generation, we systematically changed the number of total participants (*N* = 200, 600, 1,000), the number of time points (*T* = 4, 6, 8), and the size of autoregressive parameters (*β* = *β*_*x*_ = *β*_*y*_ = 0.5, 0.7) and cross-lagged parameters (*γ* = *γ*_*x*_ = *γ*_*y*_ = 0.0, 0.1, 0.2). Other parameters were the same as in the previous simulation.

We generated simulated data (100 trials for each combination) by crossing these factors, resulting in 162 (= 3(*N*) × 3(*T*) × 2(*β*) × 3(*γ*) × 3(*ψ*^2^)) combinations of factors for each pair of data generation model and data analysis model. Each simulated dataset was analyzed by the three types of analysis models. In this simulation, when improper solutions (e.g., out-of-range parameter estimates or a singular approximate Hessian matrix) were observed, the results were discarded and the simulations were repeated until the total number of successful trials was 100 for each condition. The whole simulation procedure, including data generation and analysis, was conducted in R (R Core Team^19^) using the lavaan (Rosseel^20^) package with the ML estimation method. Simulation code is available in the Online Supporting Materials.

From the results of the previous simulation, we expected a large proportion of improper solutions when applying the RI-CLPM and the STARTS model (especially when the analysis model was misspecified), which would indicate that the parameter estimates in these models might be substantially biased by discarding results with improper solutions. Therefore, we limited our attention here mainly to the differences in the standard errors of the cross-lagged parameters estimates between models. This is because the standard errors should be less influenced by the occurrence of improper solutions, given that improper solutions are mainly caused by the magnitude of point estimates (e.g., out-of-range parameter estimates or a singular approximate Hessian matrix) rather than the magnitudes of associated standard errors.

Table 4 presents the marginal means of estimated standard errors for different data generation models and analysis models under the different conditions of *N* and *T*. Note that we aggregated the other factors (*β*, *γ*, and *ψ*^2^) because we did not observe any notable influences of these factors on the estimates of standard errors. From Table 4, as we have observed from the five case studies, standard errors in the RI-CLPM and the STARTS model tend to be larger than those in the CLPM in most cases. Specifically, the standard errors were 1.1– –2.2 times the size of the CLPM in the RI-CLPM and 0.8– –4.2 times the size in the STARTS model. In applying the CLPM and the RI-CLPM, the standard errors decrease as *T* increases for correctly specified analysis models.

However, this was not the case when applying the STARTS model. Although it shows similar magnitudes of standard errors under different conditions of *T* when the analysis model was correctly specified, we saw the opposite pattern when the analysis model was incorrectly specified: the standard errors became *larger* as *T* became larger. When the true model is either the RI-CLPM or the STARTS model, standard errors with the CLPM tend to be smaller than those with other models, indicating that (incorrectly) applying the CLPM without comparing alternative models entails a greater risk of committing a type-1 error when statistically testing for reciprocal relations.

Table 5 shows the marginal means of the proportions of models reaching inconsistent conclusions about the statistical significance of cross-lagged estimates when the true value of *γ* is (a) zero or (b) nonzero. For example, in a pair of the CLPM and the RI-CLPM, the models suggested different conclusions in two ways, with the CLPM showing significant results but the RI-CLPM not, and vice versa. In each condition, the upper row counts the sum of these two proportions, while the lower row counts only the first case, where the CLPM shows significant results but the RI-CLPM does not.

From Table 5(a), it is very obvious that different models tend to show inconsistent results (in terms of statistical significance) for cross-lagged estimates. Notably, when they show different results, in most cases only the simpler model (the CLPM being compared with the RI-CLPM and the STARTS model; the RI-CLPM being compared with the the STARTS model) showed a significant result. Note that the influences of *T* and *N* vary depending on the data generation models and analysis models. However, from Table 5(b), when *γ*=0 models tend to converge to agreement more frequently, although increasing *T* and *N* increased the risk of different statistical conclusions between models.

Although we have to take care about possible biased results here as a consequence of discarding the results when improper solutions were produced when applying the RI-CLPM and the STARTS model, this simulation clearly demonstrates that statistical tests of cross-lagged effects can often show substantially inconsistent results, regardless of the number of participants or time points, especially when cross-lagged relations are actually present. One primary source of this should be the inflated standard errors of cross-lagged parameter estimates, as observed earlier.

## General Discussion

In this manuscript, we discussed limitations of the commonly-used CLPM (specifically, the conflation of between-person and within-person effects) and the importance of considering alternatives such as the RI-CLPM and the STARTS model when investigating reciprocal effects within individuals. Through a literature search, case studies, and statistical simulations, we showed the current predominance of the CLPM for testing cross-lagged effects in the medical literature and demonstrated the risk of drawing inconsistent conclusions depending on the model tested. In addition, we showed the potential risk of improper solutions when applying alternative models (the STARTS model, in particular) with the ML method, especially when the model is misspecified.

One important observation was that many researchers implicitly precluded the option of using RI-CLPM or the STARTS model by collecting data from only two time points. Given the substantially different results obtained from different models, we recommend that applied researchers collect longitudinal data at more than two time points whenever possible. If we were to assume the instability of parameters across time points, more than three time points are required to compare model fits between RI-CLPM and the STARTS model. If collecting data from a larger number of time points, then performing model selection based on model fit indices is an important step in minimizing the risk of drawing erroneous conclusions about reciprocal effects. Parameter estimation may be a serious obstacle, though, especially when applying the STARTS model. Although improving research design (e.g., by choosing an appropriate sample size) is important, choosing a different estimation strategy, such as Bayesian estimation (Lüdtke, Robitzsch, & Wagner^16^), and choosing a better specified analysis model via model selection seems to be more useful. Future research should more intensively investigate the utility of Bayesian estimation in applying various cross-lagged models.

Some limitations should be noted. First, the RI-CLPM and the STARTS model assume that autoregressive and cross-lagged parameters are fixed across participants, but we could incorporate random slopes for these effects. This would allow investigating the possible individual differences in within-person reciprocal effects. Such a model can be easily implemented with a multilevel modeling framework (e.g., Bringmann et al^21^; Schuurman, Ferrer, de Boer-Sonnenschein, & Hamaker^22^). We suspect that such new models may be more susceptible to improper solutions given the increased number of parameters and complicated covariance structure. Future investigations should provide clearer insights into how researchers can choose the appropriate analysis model in practice. A second point relates to the extension of the current discussion to other statistical models. For example, medical researchers are often interested in testing mediation effects to understand the mechanism by which one variable influences another (e.g., Richiardi, Bellocco, & Zugna^23^; Ten Have & Joffe^24^; VanderWeele^25^), and they are often assessed in a longitudinal design (e.g., Huang & Yuan^26^; Preacher^27^). The issue of the current paper applies especially to longitudinal mediation models that include cross-lagged relationships (e.g., a dynamic autoregressive mediation model; Maxwell, Cole & Mitchell^28^). If researchers fail to account for stable individual differences, then the estimated mediation effects conflate between-person and within-person processes. The current discussion is useful for considering possible alternatives when evaluating longitudinal mediation effects, and investigating the statistical properties of estimates and the frequency of estimation problems should be intriguing topics for future research. Finally, although the current study focused only on the medical literature, future study should examine common practices for testing reciprocal effects in other fields. This would give us more empiricalinsights into the similarities and differences in these cross-lagged models.

## Data Availability

All analysis and simulation codes used during this study are included in this published article (and its Supplementary Information files). The some numerical datasets analysed during the current study are not publicly available due to the data sharing policy of their sources, these datasets are however available from the authors upon reasonable request and with permission of each third party.

## Ethical Approval and Informed Consent

We have not carried out any experiments during this study.

## Additional Information (competing interests)

The authors declare no competing interests.

